# Deep learning accurately quantifies plasma cell percentages on CD138-stained bone marrow samples

**DOI:** 10.1101/2022.02.18.481050

**Authors:** Fred Fu, Angela Guenther, Ali Sakhdari, Trevor D. McKee, Daniel Xia

**Affiliations:** STTARR Innovation Centre, University Health Network, Toronto, ON, Canada; Division of Hematopathology and Transfusion Medicine, University Health Network, Toronto, ON, Canada; Scarborough Health Network, Toronto, ON, Canada; Deciphex Inc., Dublin, Ireland

## Abstract

**Background:** The diagnosis of plasma cell neoplasms requires accurate, and ideally precise, percentages. This plasma cell percentage is often determined by visual estimation of CD138-stained bone marrow biopsies and clot sections. While not necessarily inaccurate, estimates are by definition imprecise. For this study, we hypothesized that deep learning can be used to improve precision.

**Methods:** We trained a semantic segmentation-based convolutional neural network (CNN) using annotations of CD138+ and CD138− cells provided by one pathologist on small image patches of bone marrow and validated the CNN on an independent test set of images patches using annotations from two pathologists and a non-deep-learning commercial software. Once satisfied with performance, we scaled-up the CNN to evaluate whole slide images (WSIs), and deployed the system as a workflow friendly web application to measure plasma cell percentages using snapshots taken from microscope cameras.

**Results:** On validation image patches, we found that the intraclass correlation coefficients for plasma cell percentages between the CNN and pathologist #1, a non-deep learning commercial software and pathologist #1, and pathologists #1 and #2 were 0.975, 0.892, and 0.994, respectively. The overall results show that CNN labels were almost as accurate pathologist labels at a cell-by-cell level. On WSIs from 10 clinical cases, the CNN continued to perform well, and identified two cases where the sign-out pathologist overestimated plasma cell percentages.

**Conclusions:** The high labeling accuracy of the CNN supports its eventually application as a computational second-opinion tool for the measurement of plasma cell percentages in clinical practice.

## Background

Plasma cell percentage in the bone marrow is calculated by dividing the number of plasma cells by the total number of nucleated cells in a sample. Accurate percentages are critical for the diagnoses of plasma cell neoplasms [1], and for disease follow-up post-treatment. Pathologists routinely obtain these percentages by examining CD138-stained slides using one of two methods. The first is visual estimation across entire bone marrow specimens at low magnification. The second is to count cells one at a time in small selected patches of marrow at higher magnification.

Several factors challenge the accuracy and precision of human-based percentages. These include variable sizes and non-random distributions of neoplastic plasma cells. As expected, visual estimates are by definition imprecise. However, some studies have shown that pathologists may also overestimate plasma cell percentages at low magnifications when compared to corresponding cell-by-cell counts at higher magnifications --suggesting that these estimates can also be inaccurate [2]. While cell-by-cell counts are accurate and precise for small selected patches, it is difficult to be certain if these small samples are faithful representations of the much larger bone marrow specimens.

Imprecision and inaccuracy are most problematic when the true percentages of clonal plasma cells are close to an important diagnostic cut-off. In a hypothetical example, if the true percentage is 11%, many pathologists would have a difficult time determining if this is above or below the 10% threshold [1] used to separate monoclonal gammopathy of undetermined significance (<10%, a precancerous lesion) from plasma cell myeloma (≥ 10%, an often aggressive cancer).

In theory, accurate and efficient computational enumeration of CD138+ and CD138− cells across an entire slide offers a potential solution. A review of published literature, however, identified a number of limitations in existing implementations [3–8]. First, most computational studies used visual estimates as reference standards. Per Aeffner et al. [9], this creates a “gold-standard paradox” where algorithms are trained and optimized using potentially unreliable references, subject to visual biases, which computational methods were supposed to avoid. As a result of the focus on whole slide estimates, most studies also did not report on concordance between algorithms and humans at a cell-by-cell level. Finally, most studies relied on older area-based and cell- or nuclear-segmentation-based methods, and until very recently [10], did not employ deep learning.

Accordingly, our solution was to start at the cell-by-cell level using small patches of marrow. In principle, this addresses issues such as non-random distributions and variable sizes of tumor plasma cells that plague visual estimates, and avoids the “gold-standard paradox.” Specifically, we trained a convolutional neural network (CNN) using raw pixel data and labels of CD138+ and CD138− cells provided by one pathologist, without reference to prior cell- and nuclear-segmentation. Once trained, the CNN was benchmarked against a commercially available “classical computer vision” (non-deep-learning) software and a second pathologist on an independent set of small image patches, using new labels from the original pathologist as reference. Once satisfied with the performance of the CNN, we scaled up our system to whole slide images (WSIs), and deployed it as a workflow-friendly web application.

## Methods

### Case collection, CD138 immunohistochemistry, pathology report data, and ethics statement

16 cases with bone marrow biopsies and/or clot sections were selected from the archives of the Department of Pathology at the University Health Network (Toronto, Ontario, Canada). These include cases of plasma cell neoplasms (n = 7), myeloid neoplasms with reactive plasma cells (n = 1), and normal bone marrows with reactive plasma cells (n = 8). Anti-CD138 antibody was used for detection of plasma cells by immunohistochemistry (MI15 clone; Agilent Dako, Santa Clara, California, USA) at 1:25-1:50 dilution range.

Plasma cell percentages for 10 clinical pathology reports were obtained from institutional pathology laboratory information management system. For plasma cell percentages with range estimates, middle values were used for analyses (e.g., 5-10% was recorded as 7.5%). Plasma cell percentages reported as “normal,” “not increased,” or “less than 5%” were recorded as 2.5% (middle value of 0-5%) for analyses.

The study is authorized by the UHN Research Ethics Board (CAPCR 18-5069).

### Patch selection and annotation

Slides were scanned at 40x magnification (0.25 μm/pixel resolution) using an Aperio AT2 scanner (Leica Biosystems, Wetzlar, Germany) and imported into the open-source digital pathology platform QuPath [11] for image patch selection and annotation. In each whole slide image, tissue was separated from background and the resulting tissue area was divided into 512 by 512 pixel tiles. From each slide, representative tiles were selected for annotation. Within the selected tile, one pathologist labeled the centre of each nucleated cell marking it as CD138+ or CD138−. Anucleate red blood cells, megakaryocytes, endothelial and stromal cells were not labeled. If a cell was only partially within the tile but its centre was not within the tile boundaries, the pathologist was instructed to ignore it. A customized script was used to export these tiles with an additional 32 pixels of padding on each side which is retained for context (resulting in a final size of 576 × 576 pixels). Similarly, a set of encoded “label” images were exported as 8-bit images of equal size in which pixels corresponding to negative or positive cell annotations were assigned numeric values of 1 or 2, for negative or positive cell annotations respectively, and otherwise 0 for background.

101 tiles (each consisting of an image and labels) from the biopsy sections of 5 subjects constituted the training (80%) and validation (20%) datasets used for model tuning and selection. Another 20 tiles from 10 additional subjects constituted a held-out test set which was solely used for evaluation on the final model. These test set slides were also scanned at a separate microscopy facility in order to better assess generalizability across scanning equipment.

### Deep learning-based analysis and model training

This deep learning approach utilizes a convolutional neural network in a encoder-decoder or U-Net-like configuration [12]. Briefly, this network architecture involves a series of “encoder” layers which maps input images into high-dimensional feature vectors, and a subsequent series of “decoder” layers which maps these feature vectors back into pixel-wise classifications, allowing the network to be used for semantic segmentation (the classification of individual pixels of an image). Here, the encoder network used is VGG-based [13] with batch normalization layers. To avoid needing to learn basic features from scratch and to reduce training time, transfer learning was employed by initializing the encoder network with pre-trained weights from ImageNet. For the decoder network, up-sampling is performed using bilinear interpolation combined with convolutional layers. Note that as this network has a fully convolutional architecture [14], inference is only limited by the size of available VRAM, allowing for images of various sizes to be evaluated.

Input images were normalized and labels were also preprocessed using a grayscale dilation operation to provide additional context surrounding the annotated points. The training dataset was also augmented by employing random transformations such as flips, rotations (with subsequent reflect padding to avoid blank image regions), and brightness adjustments in order to increase the size of the dataset and improve generalizability.

Training was conducted on the training samples using stochastic gradient descent on the categorical cross-entropy loss function on individual hyperparameter configurations for up to 200 epochs. Hyperparameters included learning rate and class weighting (i.e. relative importance of negative cells, positive cells, and background), which were optimized using a grid search approach.

A 2D softmax function was used to convert the model outputs into two pseudoprobability heatmaps (one for CD138− cells and one for CD138+ cells). These heatmaps were post-processed using 3×3 median and Gaussian filters and a final peak detection filter was used with a minimum distance constraint of 10 pixels to identify the centroids of CD138− and CD138+ cells. A linear sum assignment algorithm was used to match these point detections against the ground truth labels based on their proximities so that each detection could be placed in the three-class (CD138+, CD138−, and background) confusion matrix. Pairs were considered matching if they were within a distance threshold of 15px (3.75 µm), similar to other approaches [15,16].

The final model that was selected was the one that attained the best macro-averaged F1-score [17] on the validation set, computed based on the confusion matrix during training. We found that incorporating padding around tiles during the evaluation improved the network’s performance by giving it additional context from outside of the annotated region (e.g. if a nucleus was split in half at the border). Macro-averaged F1-score (a balanced metric) was used since it treats the detection of positive and negative classes equally, which is important as the diagnostic criteria employs a ratio of CD138-positive and CD138-negative cell counts.

Model training and inference was carried out on an NVIDIA Titan Xp with 12 GB of VRAM. The network was implemented in Python 3.7, notably using the PyTorch [18] (version 1.2.0) to implement the neural network and for pre-trained VGG weights, and scikit-image (version 0.15.0) [19] for post-processing.

### Analysis with commercial digital pathology platform

To have as a basis for comparison, a proprietary algorithm from the digital pathology software Definiens TissueStudio (version 4.4.2) was employed, which uses either area-based and cell segmentation-based analysis in a similar general manner to previous literature reports for computational plasma cell counting. In both cases, staining parameters were configured to detect DAB membrane staining.

The “nucleus detection” and “membranes and cells” algorithms were used to perform cell-based segmentations, which use color deconvolution to separate hematoxylin and DAB staining to identify nuclei and membranes, respectively. The parameter values and thresholds for these algorithms were fine-tuned by visual inspection amongst patches from specific images in the training set. The centroids of the CD138+ and CD138− cell segmentations were then extracted as detections for counting. Comparison against ground truth labels was performed using the same matching algorithm as described for the CNN.

### Whole slide inference

Once the CNN model was trained, it could be applied to analyze whole slide images (WSIs). At full resolution, WSIs are typically on the order of gigapixel images and are too large to pass through the model, requiring specialized handling. Here, a “shift-and-stitch” approach [20] was employed, where a whole slide Aperio SVS file was loaded in patches using the VIPS library [21] by running a sliding window (2048 × 2048 pixels) over the full image and analyzing each patch sequentially. These patches were “padded” on each side by an additional 32 pixels to retain context at edges, resulting in a 2112 × 2112 pixel tile. At the edges of each WSI, where padding outside of the image was required, reflection padding was used. Optionally, a tissue mask annotation image created using QuPath could be supplied, which reduced computational burden by ignoring unmasked regions from being processed by the model pipeline. Finally, for easy visualization of the results on WSIs, the final point detections were outputted into a QuPath-compatible text file which could be loaded into QuPath as point annotations back onto the original WSIs.

### Statistical methods

To report performance across the entire testing dataset, F1-score (mentioned previously for evaluating performance on the validation set) and ICC were used. F1-score was derived from the resulting confusion matrices across all predicted detections and actual annotations from applying the model to the test dataset. ICC measures the agreement across groups. Intraclass correlation coefficient (ICC) [22] was calculated using the irr package in R (version 3.6.2), specifying a two-way model and absolute agreement. Both methods can be applied to make comparisons between two “raters” (either a human annotator or a model-based detection).

## Results

### Plasma cell percentage determination using commercially-available software

Prior to expending resources to develop a CNN, we first explored quantification of CD138+ and CD138− cells using a non-deep learning commercial software: Definiens Tissue Studio. We found that the “nucleus detection” and “membranes and cells” algorithms from the software correctly labeled the majority of plasma cells and hematopoietic cells (Figure 1). However, there were also occasional labeling errors. Of note, in areas with high non-specific staining, the software incorrectly segmented additional “plasma cells” (Figure 1A and B; indicated by black arrows), which illustrates a potential drawback of relying on segmentation-based methods, notably, the occurrence of over-segmentation (a common observation for computational cell segmentation). Since our goal was to eventually deploy a software tool to assist hematopathologists with clinical diagnostics, we felt strongly that labeling errors should be minimized as much as possible. As such, we turned to deep learning to evaluate it as a potentially more accurate solution (Figure 1C).

**Figure 1.**
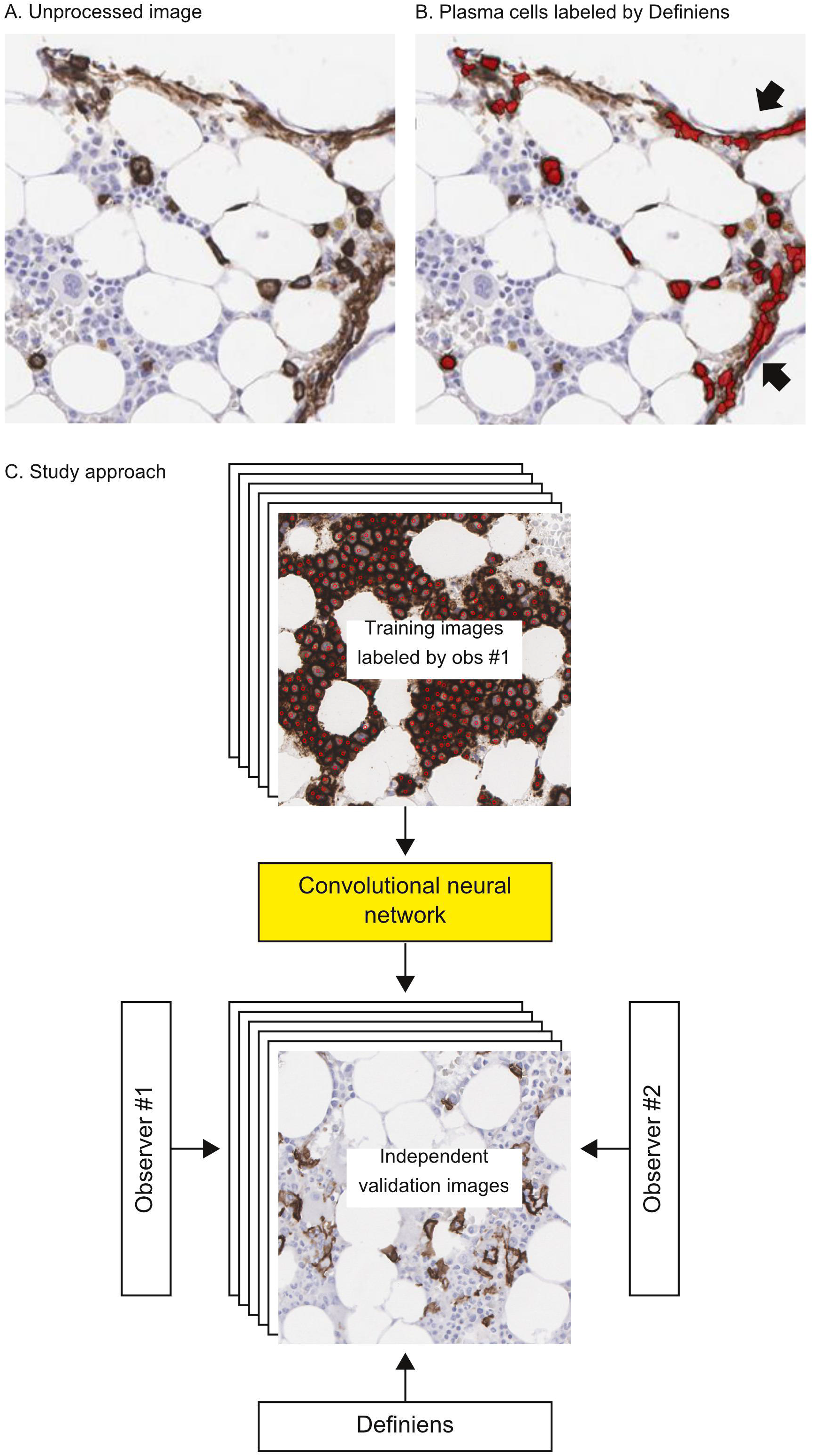
Overall approach. Initially, we used a commercial software (Definiens Tissue Studio) to count CD138+ and CD138− cells. A-B shows that many plasma cells were correctly identified, but the software also made mistakes, for example, labeling edges of tissue with non-specific staining as plasma cells (indicated by arrows). To us, this example served as a cautionary tale for relying on segmentation. C) Hypothesizing that a deep-learning approach could perform better, we trained a CNN using small patches of images of bone marrows labeled for CD138+ and CD138− cells by a pathologist (observer #1; only positive labels are shown in the figure). The CNN was then tested on an independent set of validation image patches. The CNN’s labels were compared against labels generated by Definiens and two observers (#1 and #2). Abbreviations: CNN – convolutional neural network.

### Model training

Since it is controversial whether low magnification visual estimates of plasma cell percentages are appropriate gold standards, we trained our model using point-by-point annotations of CD138+ and CD138− cells provided by a pathologist across small image patches of bone marrow biopsies and clot sections. Counting the number of CD138+ and CD138− cells in an image can be formulated as a multi-class detection task where the centers of cells are identified and then enumerated. This is contrasted with a segmentation task, as distinguished by Janowczyk et al [23], which tries to identify a contour around each cell. By doing this, cells only need to be labelled by marking their centroids; this is particularly suitable for plasma cell detection because it can be difficult to objectively delineate cytoplasmic boundaries between some cells. We build on work from past approaches which identify the presence of a single cell subtype [15,23] and extend it to the detection of two different cell types: CD138- and CD138+ cells. The overall idea is to optimize model performance using these small image patches and then scale-up the point-by-point labeling to whole-slide images.

### Evaluation of CNN labels against pathologist and commercial software labels on an independent validation image patch set

Following model training, we benchmarked the CNN’s labels against labels provided by two pathologists (observers #1 and 2), as well as labels generated by Definiens, on an independent set of small image patches. Figures 2A-D and 3A-D show an example of a validation patch where there was a high degree of labeling concordance across the two observers and two computational methods. By contrast, Figures 2E-H and 3E-H show the results from a more challenging validation patch. Here, it appears that the high background staining caused the commercial software to count far more plasma cells and fewer hematopoietic precursors in comparison to the CNN and the two observers.

**Figure 2.**
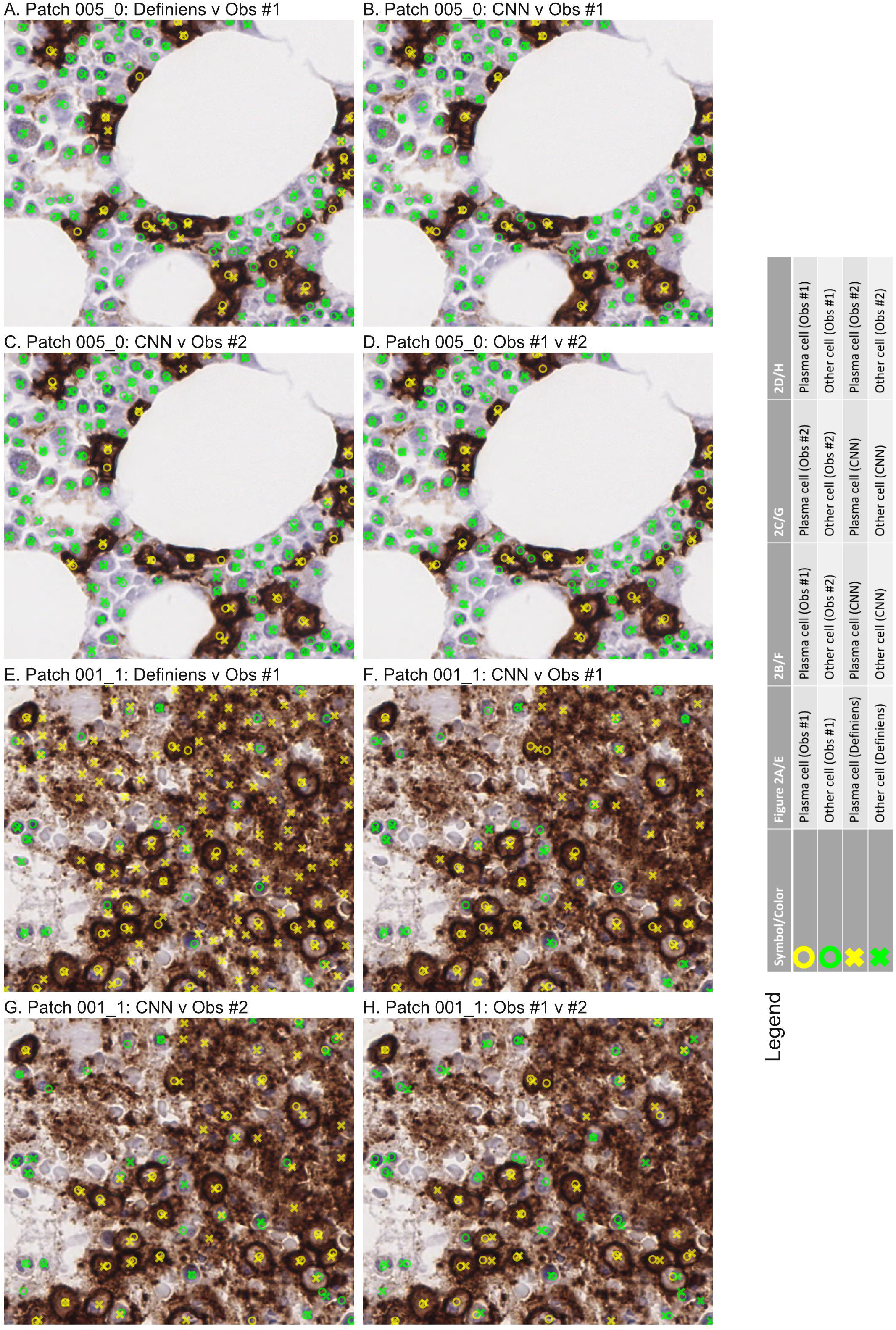
Comparison of labels from two human observers (#1 and #2), Definiens, and CNN on independent validation. This shows actual labels from A and E) Definiens v observer #1, B and F) CNN v observer #1, C and G) CNN v observer #2, and D) observer #1 v #2 for two image patches. Symbols and colors are indicated in the legend. A-D) Patch 005_0. This proved to be a relatively “easy” patch to label. The labels from the CNN, Definiens, and two observers were largely concordant. E-H) Patch 001_1. By comparison, this patch was more “challenging.” Here, the Definiens software labeled many more plasma cells than the CNN and two observers, possibly because there was high background staining, which we previously observed (e.g., Figure 1). All patches shown are 512×512 pixels corresponding to physical dimensions of 128×128 µm. Abbreviations: CNN – convolutional neural network.

**Figure 3.**
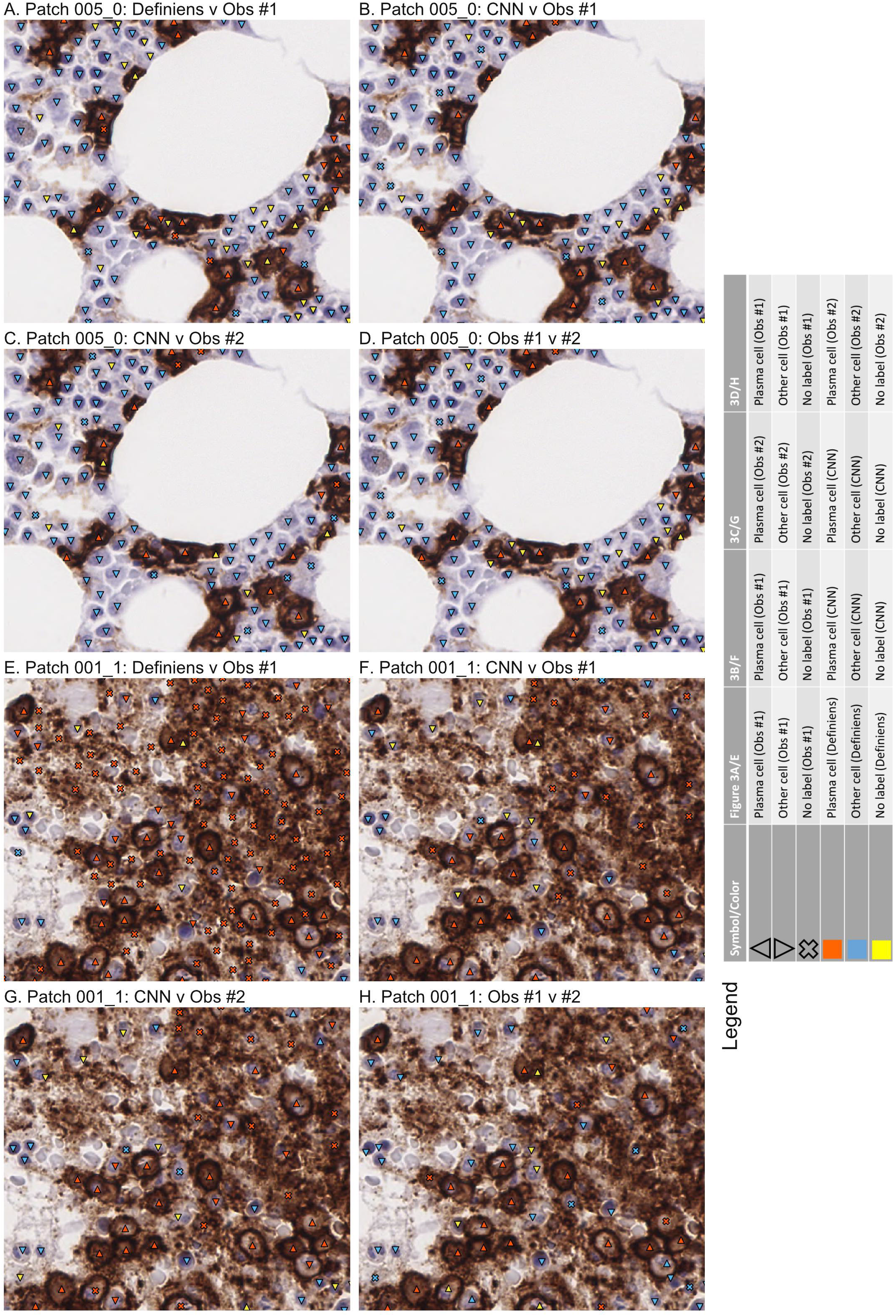
Integrated labels from two observers (#1 and #2), Definiens, and CNN. Symbols and colors are indicated in the legend. The raw image patches and order are the same as for Figure 2. All patches shown are 512×512 pixels corresponding to physical dimensions of 128×128 µm. Abbreviations: CNN – convolutional neural network.

The results across the ten validation patches are summarized in Figure 4. In general, various measures of label concordance were the lowest between the commercial software and observer #1 (Figure 4A), and highest for the two human observers (Figure 4D). Using observer #1’s labels as reference, the CNN performed significantly better than the commercial software, and slightly worse when compared to observer #2. Intraclass correlation coefficients for plasma cell percentages between the CNN and pathologist #1, a non-deep learning commercial software and pathologist #1, and pathologists #1 and #2 were 0.975, 0.892, and 0.994, respectively. Overall, these data suggest that the CNN achieved a level of concordance that approaches but is slightly inferior to pathologist-based point-by-point annotations.

**Figure 4.**
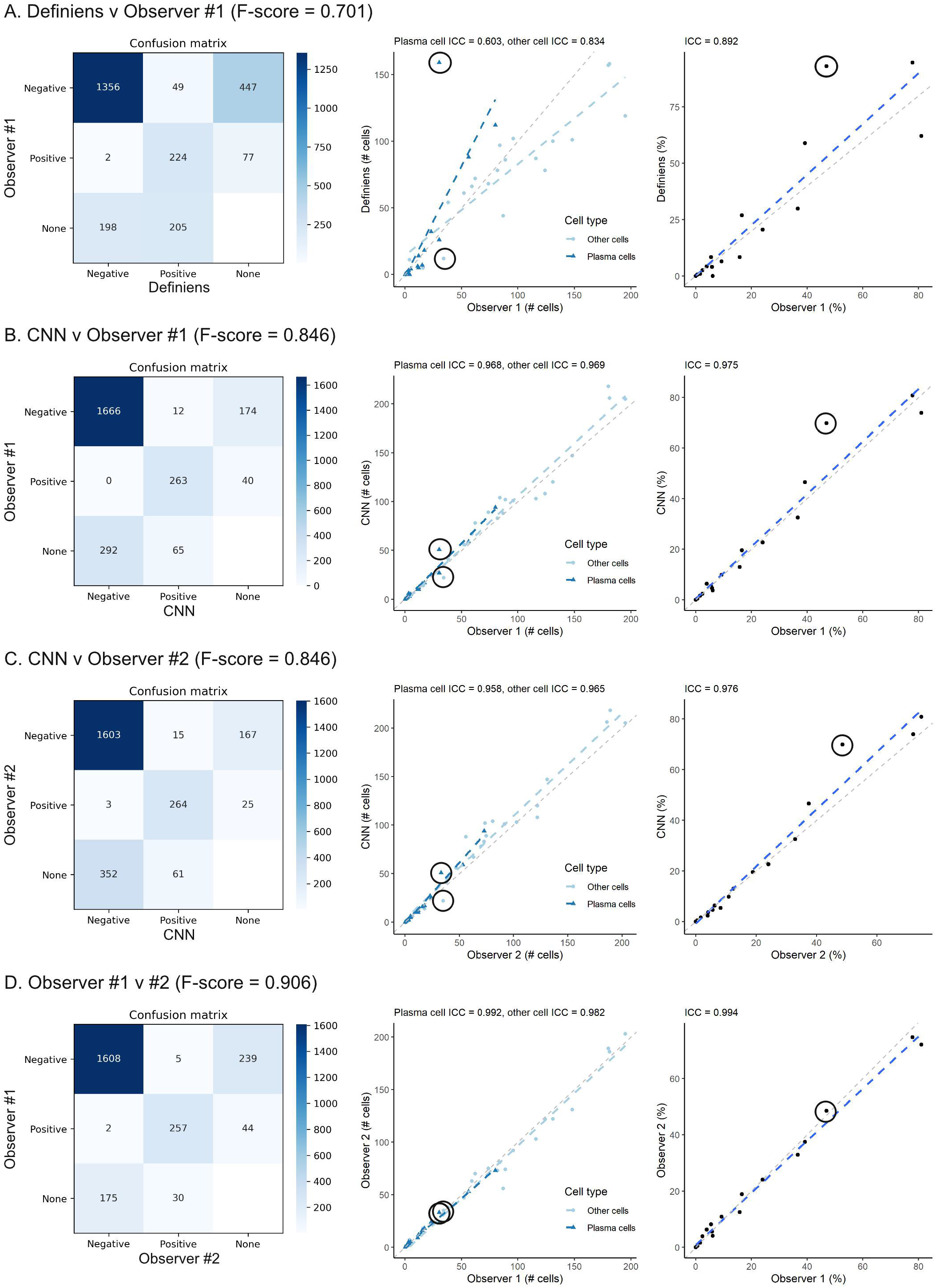
Summary of validation on small image patches. Confusion matrices with F-scores for cell-type labels (left), scatterplots for cell-type labels (mid), and scatterplots plasma cell percentages (right) are provided for four comparisons: A) Definiens v observer #1, B) CNN v observer #1, C) CNN v observer #2, and D) Observer #1 v #2. Results from the challenging image patch (001_1; Figures 2E-H and 3E-H) is indicated with circles. ICCs are indicated at the top of each scatterplot. Using observer #1 as reference, the CNN performed significantly better than the commercial software, and approached the level of labeling accuracy for pathologist #2. Abbreviations: CNN – convolutional neural network; ICC - intraclass correlation coefficient

### Whole Slide Image analysis

Since counting cells one-at-a-time across an entire biopsy would be impractical for hematopathologists, a true test of the potential utility of the CNN would be to assess its point-by-point labels using real-size bone marrow specimens. For this, we applied the CNN to pathologist-annotated regions of interest across 10 whole-slide images (WSIs) of bone marrow biopsies and clot sections, and compared the resulting plasma cell percentages against those from the corresponding bone marrow pathology clinical reports.

Figure 5A-B shows that there was near-perfect concordance between CNN predictions and pathology reports for 8 out of 10 WSIs. For the two remaining cases, the CNN predicted a lower percentage, while pathologist(s) reported a higher percentage on clinical reports. Figure 5C shows a representative image from one of the two discordant samples. In this image, plasma cells appear to occupy slightly more than half of the bone marrow biopsy by area; however, by relative numbers of blue (other cell) and orange (plasma cell) labels, it is clear that the true plasma cell percentage is still less than 50%. Since the small number of labeling errors by the CNN cannot account for such a large difference in percentages, the CNN’s predicted percentage (29.3% across the entire biopsy) is more accurate than the estimate that appeared in the clinical report (about 60%).

**Figure 5.**
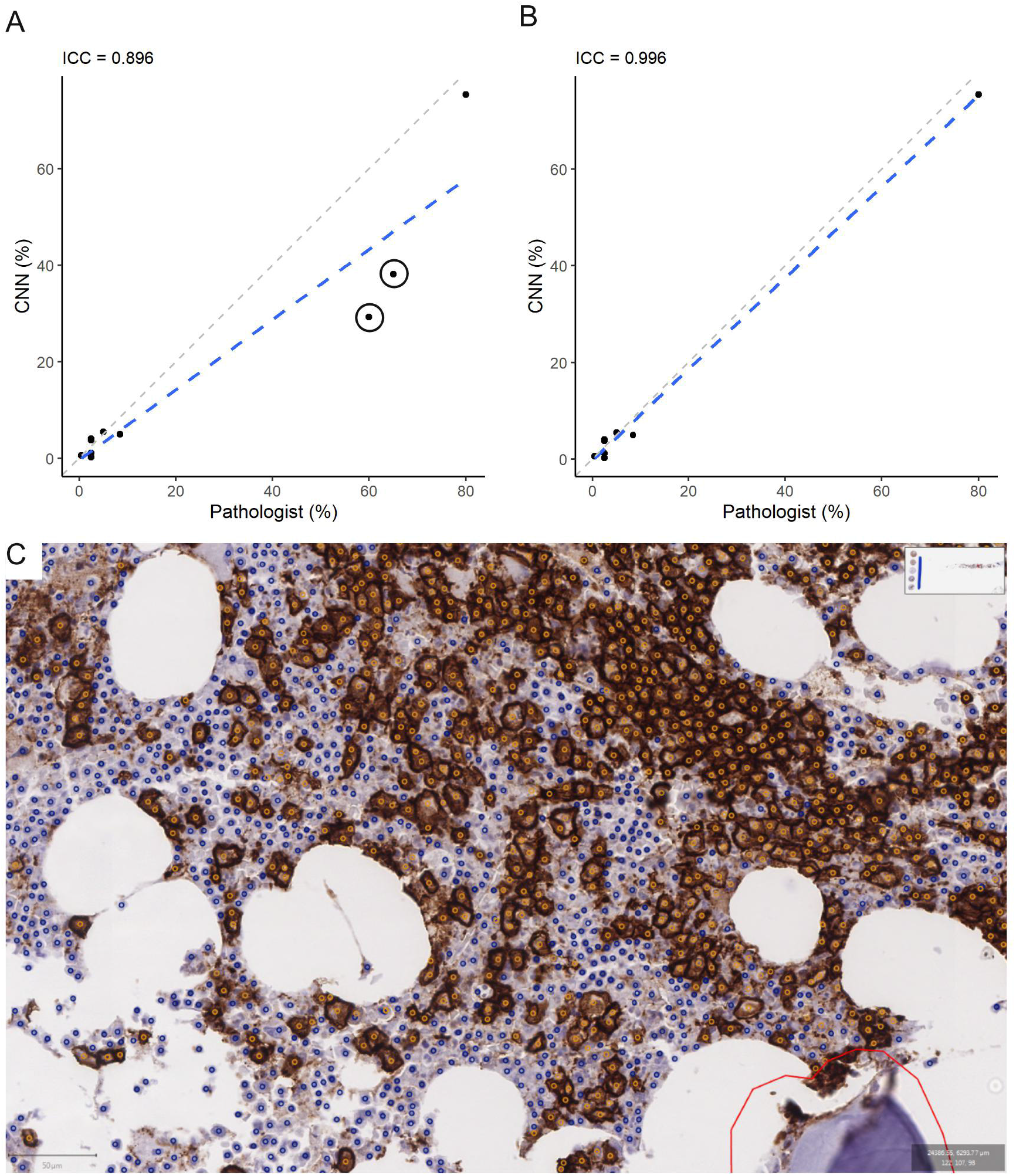
CNN labeling of plasma cells on WSIs. A) The ICC for plasma cell percentages on WSIs from CNN and from pathology reports was 0.896. There were two outlier cases where the predicted percentages from the CNN were much lower than those reported by pathologists (indicated with circles). B) If the two cases were removed, the ICC improves to 0.996. For the two problematic cases, one was a core biopsy, and the other was a clot section with many small marrow particles, each with variably abundant plasma cells. C) CNN-labeled image of the core biopsy case shows that while plasma cells (orange labels) appear to occupy slightly more than half the cellularity by area, these plasma cells are less frequent than other hematopoietic cells (blue) overall (scale bar on lower left is 50 µm in length). Since there does not appear to be significant labeling errors by the CNN, it is likely that the CNN’s percentage is more accurate and the pathologist overestimated. The red manual outline delineates the border of the region of interest within which the CNN was applied. Abbreviations: CNN – convolutional neural network; ICC – intraclass correlation coefficient; WSI – whole slide image.

A similar conclusion was drawn when the second discordant case was reviewed. This WSI was from a clot section with numerous small bone marrow particles, each with variable percentages of plasma cells. This type of sample is very difficult for pathologists to estimate. Again, the relative rarity of labeling errors on review suggests that the CNN’s prediction was more accurate (38.2%), and that the pathologist overestimated on immunohistochemistry (60-70%). The CNN’s prediction was also closer to the plasma cell percentage on the aspirate differential for this case (45%).

While the difference in plasma cell percentages would not have changed the overall diagnoses for the two cases (de novo and persistent plasma cell myeloma, respectively), the overall excellent performance for the CNN illustrates its potential as computational second-opinion tool for WSIs.

### Deployable web-based server for microscope camera snapshots

The results presented above suggest that one potential workflow would be to apply the CNN to digitally scanned WSIs of problematic CD138-stained bone marrow biopsies and clot sections. However, the scanning and analysis of WSIs adds to clinical turnaround time, which could discourage the use of this potentially helpful tool. Further, transfer of WSIs from one institution to another for analysis would involve very large files, which may be impractical to work with at high case volumes. In view of this, we developed a simple web application to facilitate more practical clinical workflows, where pathologists can capture snapshots of CD138-stained images of bone marrow samples (at 400X magnification), taken directly from microscope-mounted cameras during routine sign-out, and upload those snapshots to a server (Figure 6), to receive near-real-time plasma cell percentages for the snapshot image. The server backend was implemented using Python 3.7 and Flask to retrieve the uploaded image and run it through the analysis pipeline. The output was served using a lightweight web interface primarily using the mapping library OpenLayers to display the images and detected points. In our trial, the web application was hosted on the same workstation as was used to train the model and accessible within the institution’s internal network. The advantage of this approach is that it does not require users to have any hardware or software installations beyond an Internet browser, making it easily accessible. The application returns absolute counts of positive and negative cells, plasma cell percentages, as well as a screenshot labeling each class of cells. The detection algorithm generated results within a few seconds of image upload, supporting its integration in routine clinical workflows.

**Figure 6.**
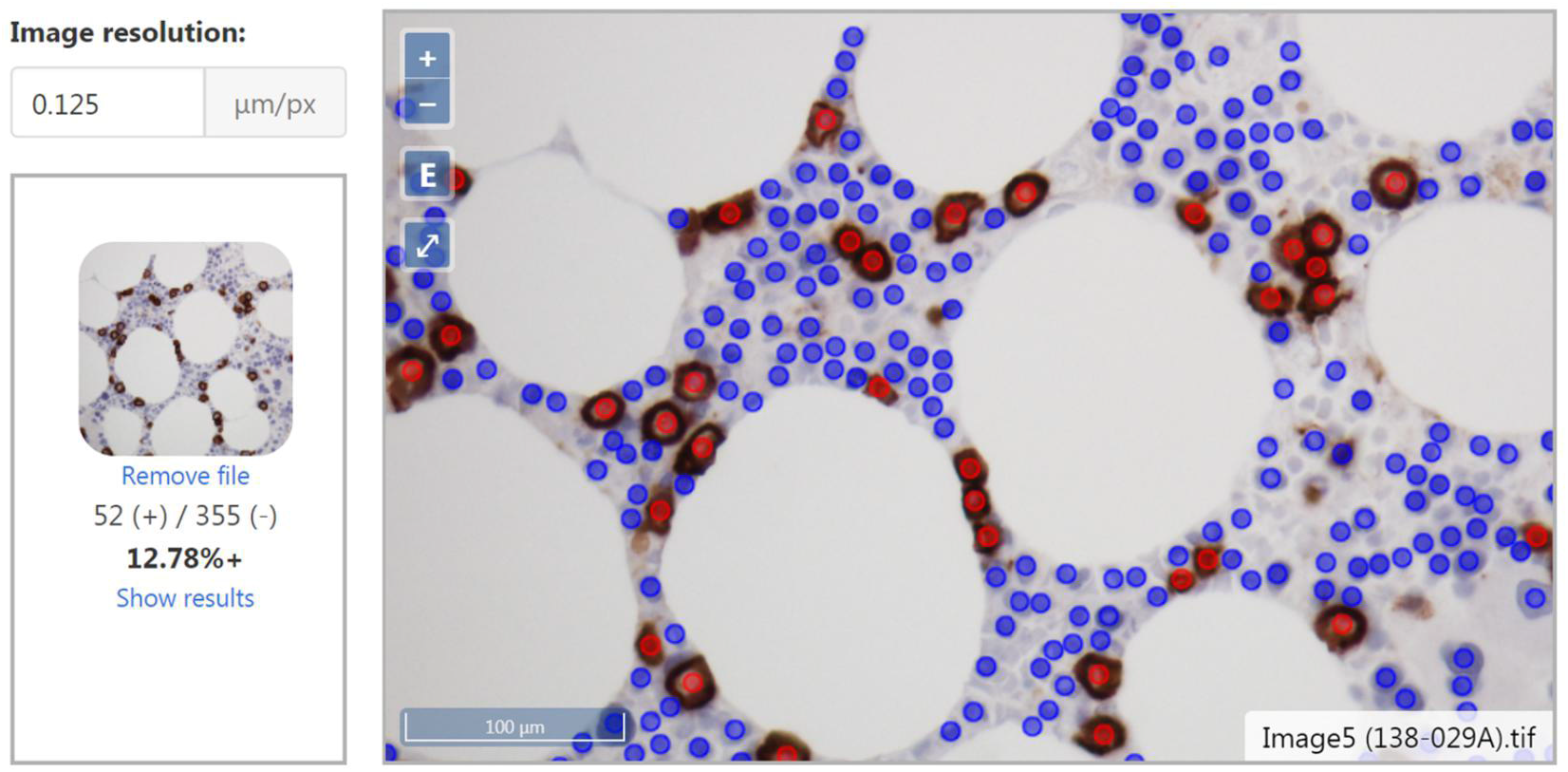
Web application user interface. The Flask application applies the CNN to an uploaded image taken from a microscope camera. An unprocessed thumbnail, numbers of positive and negative cells, and percentages are on the left. Labeled image with scale bar is shown on the right.

## Conclusions

Membranous and cytoplasmic stains can be difficult to evaluate by approaches that focus on segmenting cells and nuclei. It is possible for software to over-segment areas with intense non-specific staining into additional “cells” that do not actually exist (e.g., Figures 1, 2E-H, and 3E-H) and/or miss nuclei that are comparatively weakly-stained for hematoxylin (for which the existing hematoxylin signal is buried under the stronger DAB signal). By contrast, our CNN avoids segmentation, and is trained instead to directly predict where pathologists would place labels for cells on bone marrow images, by relying on the combination of pixel color intensities and local texture features. This difference is likely the main reason why our CNN outperformed classical segmentation methods employed by Definiens, achieved a level of labeling concordance that approached that for a second pathologist for small image patches, and continued to perform well when scaled-up to WSIs.

Our approach is therefore distinct from previous studies, which generally used less accurate area- or segmentation-based methods to identify/quantify plasma cells, and relied on low magnification visual estimates from pathologists as gold-standard references [3–8]. One exception is a recent publication from Baranova et al. [10] that also started with small image patches, before scaling up to WSIs. The major technical difference between our study and Baranova et al. [10] is that they relied on about 40 features from QuPath’s cell segmentation algorithm to train a downstream neural network, while ours is a single-stage CNN trained on raw image pixels and pathologist labels. We speculate that our CNN’s independence from prior segmentation explains why our reported measures of concordance appear to be higher than that reported by Baranova et al., although a fair comparison would involve a head-to-head evaluation using validation images from both studies.

As such, limitations of our study include the inclusion of only a small number of samples from a single institution, which do not necessarily permit generalizations regarding performance on image sets from outside sources and overall clinical utility. For our Flask web server, it is also unclear how many high magnification fields/images are required to faithfully represent the entire bone marrow biopsy/clot section, given the non-random distribution of plasma cells. While our current implementation for WSIs involves limiting the analysis to pathologist-identified regions-of-interest, future implementations would require automated identification of suitable areas of marrow for scoring, while avoiding bone and tissue artifacts (e.g., folded marrow sections).

Overall, the performance of our CNN supports its eventual application as a digital second-opinion tool for evaluating CD138-stained marrow samples in hematopathology --since CNN labels are highly concordant with pathologist labels at a cell-by-cell level. We note that a single-stage CNN to predict pathologist labels independent of prior segmentation could also be applied to other stains. As examples, CD34+ blasts and HER2+ breast cancer cells are arguably more challenging to score than CD138+ plasma cells, since these may show incomplete staining of tumor cells at variable intensities. The manners by which these stains are evaluated via human eyes are also not easily converted to fixed pre-defined parameters. As such, in comparison to segmentation-based approaches, we hypothesize that single stage CNNs trained on raw pixel data and pathologist labels may prove to be the superior solution for many cytoplasmic and membranous stains.

## Data Availability

The trained model and analysis server code are available online at [https://github.com/STTARR/plasma-cell-detection] with examples of usage included. Training data and model training code are currently only available upon request.

## Acknowledgements

This study is funded by the Academic Enrichment Fund provided by Pathology Associates at the University Health Network. The authors would like to acknowledge the Spatio-Temporal Targeting and Amplification of Radiation Response (STTARR) program and its affiliated funding agencies.

## Author Contribution Statement

D.X., T.D.M., F.F. designed the study, participated in data analysis, and wrote the manuscript. D.X., A.G. and A.S. provided annotations upon which the network was trained, and provided domain expertise. F.F. designed and implemented the CNN and web server. F.F. and D.X. generated figures and results. D.X. obtained funding to support the work. All authors participated in conceptualization and read and approved the final manuscript.

